# Quantification of DNA damage induced γH2AX focus formation via super-resolution dSTORM localization microscopy

**DOI:** 10.1101/638361

**Authors:** Dániel Varga, Hajnalka Majoros, Zsuzsanna Újfaludi, Miklós Erdélyi, Tibor Pankotai

## Abstract

In eukaryotic cells, each process, in which DNA is involved, should take place in the context of chromatin structure. DNA double-strand breaks (DSBs) are one of the most deleterious damages often leading to chromosomal rearrangement. In response to environmental stresses, cells have developed repair mechanisms to eliminate the DSBs. Upon DSB induction, several factors play roles in chromatin relaxation by catalysing the appropriate histone posttranslational modification (PTM) steps, therefore promoting the access of the repair factors to the DSBs. Among these PTMs, the phosphorylation of the histone variant H2AX at its Ser139 residue (also known as γH2AX) could be observed at the break sites. The structure of γH2AX focus has to be organized during the repair as it contributes to accessibility of specific repair proteins to the damaged site. Our aim was to develop a quantitative approach to analyse the morphology of individual repair foci by super-resolution dSTORM microscopy to gain insight into genome organization in DNA repair. We have established a specific dSTORM measurement process by developing a new analytical algorithm for gaining quantitative information about chromatin morphology and repair foci topology at individual γH2AX enriched repair focus. By this method we quantified unique repair foci to show the average distribution of γH2AX clusters. By monitoring γH2AX signal, we could reach 20 nm spatial resolution and resolve a single DNA damage spot, which allow us to identify different chromatin sub-clusters around the break site. Additionally, based on our new analysis method, we were able to show the number of nucleosomes in each sub-cluster that could allow us to define the possible chromatin structure and the nucleosome density around the break sites. This method is the first demonstration of a single-cell based quantitative measurement of a discrete repair focus, which could provide new opportunities to categorize spatial organization of dot patterns by parametric determination of topological similarity.

## INTRODUCTION

The DNA in the nucleus is constantly targeted by different damaging agents deriving from both endogenous and exogenous sources. DNA double-strand breaks (DSBs) are the most deleterious lesions, therefore they have to be repaired as quickly and efficiently as it is possible to prevent chromosomal loss and translocation. Since DSBs affect DNA integrity simultaneously with the recruitment of early DNA repair factors, a DNA damage response (DDR) is activated in the cells, which can arrest the cell cycle ^1,2^. For efficient DDR activation, different DSB sensors are required to activate chromatin reorganization and recruitment of downstream repair proteins that can eventually accomplish the efficient repair process ^3^. Recent studies have already shown that DNA damage can lead to immediate chromatin relaxation around the site of the damage ^4–6^. One of the first steps of DSB induced chromatin reorganization is the phosphorylation of the histone variant H2AX at its S139 residue (called γH2AX) in the proximity of the damaged site ^4,5,7,8^. The γH2AX enriched chromosomal locus, considered as repair focus, marks the damage site to initiate the recruitment of further repair proteins required for the process and performance of the repair ^9,10^. Several factors, such as cell cycle state, functional activity of genes, break position along the DNA sequence, temporal state of DNA compaction, number of simultaneously occurring DSBs, etc. have been known to influence this process, thereby assigning the fate of the cell ^11,12^. The γH2AX signal detection is regularly used to visualize and quantify the extent of DSBs and to follow the DNA repair kinetics. Several techniques have been applied to follow the changes in γH2AX signal intensity. Chromatin immunoprecipitation studies have revealed that the γH2AX signal shows asymmetrical distribution around the damage site with lower density at the transcribed regions ^13–20^. It has been already shown that both the genome topology and chromatin state are crucial for the organization of the recruitment of repair proteins. A recent study based on chromosome conformation capture experiments has highlighted the complexity of genome re-organization, including megabase range associations between certain chromosomal regions as well as smaller genomic interactions, which involve kilobase-length DNA segments ^21,22^. Additionally, optical methods, such as conventional confocal microscopy have also been regularly used for mapping the spatial distribution of DSBs. Due to the spatial resolution of these methods, the DDR signal can be detected in maximum 300 nm resolution. By using these techniques, it was shown that the γH2AX signal distributes up to a megabase around the damaged site ^8^ generating DNA repair foci with a typical feature size of half a micron, which is just above the resolution limit of traditional fluorescence microscopes. High resolution imaging based datasets of DSB structures have already been published and demonstrated via single molecule localization methods (SMLM) ^23–28^, structured illumination microscopy (SIM) ^26,29,30^, and stimulated emission depletion (STED) ^29–31^ super-resolution methods. These images can be further evaluated by using cluster analysis, and both the spatial distribution and the geometrical parameters of foci and even nano-foci can be determined ^32,33^. Single molecule localization methods (SMLM), such as dSTORM, provide the highest spatial resolution among optical methods and open the way for imaging biological structures in the sub-20 nm regime ^34–36^. SMLM determines the positions of individual molecules, which are used to create the final image ^37–39^. Such an image registration and processing method is especially appropriate for quantitative evaluation. Although dSTORM separates the sub-domains of the repair focus, quantitative evaluation of the images has been still challenging because the number of detected localizations (*N*_*localizations*_) generated by a single-labelled histone has been still unknown. Therefore, for quantitative analysis, the number of localizations per labelled histone has to be statistically determined.

Here we provide insight into γH2AX distribution in nanometre resolution by using super-resolution dSTORM microscopy technique applied either on U2OS cells exposed to neocarzinostatin treatment or on AsiSI endonuclease-expressing DIvA cells ^18,40^. By surpassing the limitation of classical confocal microscopy, super-resolution dSTORM microscopy possesses high prospecting capacity, which allows us to enlarge complex structures at γH2AX-covered chromatin regions in 20 nm resolution. With this technique we measured the DDR profiles of several genomic regions and gained temporal, functional and structural insights into the damaged chromatin units evolved during DSB repair. By means of dSTORM, in accord with already published data, we observed that the number of γH2AX foci increased following DSB induction ^41,42^. In addition, we demonstrated that the sizes of these foci were extended under DSB formation and we could also provide a higher resolution of the foci spatial organization using our new statistical approaches. This article preconcerts the nano-scale organization of the repair foci, which could highlight the spatial localization of the sub-domain structure and quantitative measurements of these repair centres.

## RESULTS

### The experimental system and the determination of the parameters used in dSTORM

The typical size of a DNA repair focus is about half micron, just above the resolution limit of traditional fluorescence microscopes. However, at high density of the DNA breaks when individual foci merge and form larger blobs the traditional imaging methods cannot be utilized. In such cases optical super-resolution microscopy is required to distinguish individual foci and reveal their sub-structures containing 20-60 nm nano-foci ^43^. In order to prove this, we generated DNA DSBs by applying neocarzinostatin and 4-OHT treatment to U2OS and DIvA cells, respectively ^14,40^. We quantified the DSB triggered γH2AX foci formation 2 hours following the break induction by labelling H2AX S139 phosphorylated sites with fluorophore-conjugated antibody. We generated traditional EPI fluorescent and high resolution dSTORM images from nuclei of non-treated and treated cells (Fig 1 A vs B). By performing dSTORM, we observed an increase in the number and distribution of γH2AX foci following DSB induction. In higher magnification of individual foci, we could identify sub-structures that we used in our quantitative analysis (Fig 1 C).

**Figure 1.**
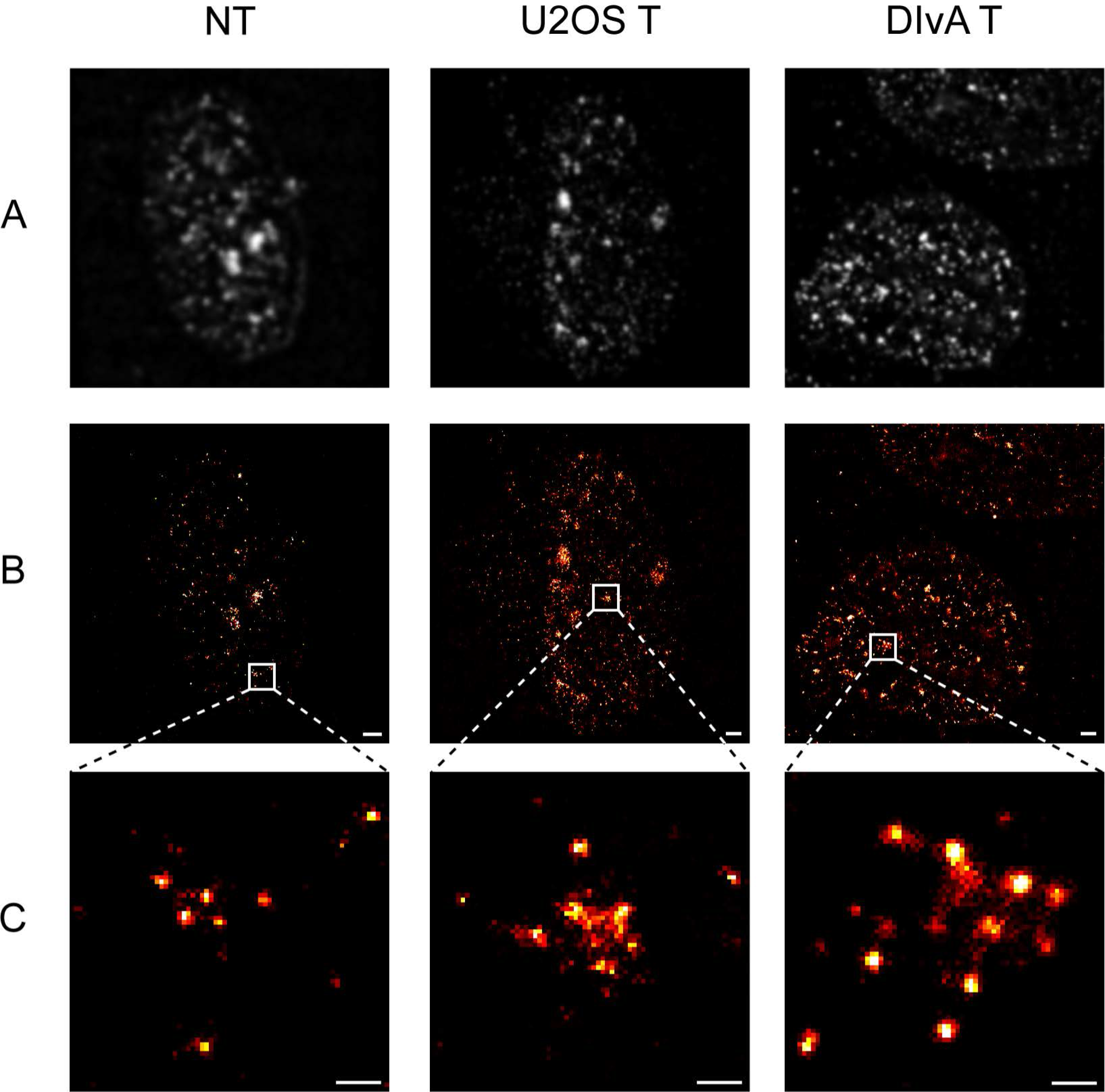
Traditional EPI fluorescence (A) and dSTORM super-resolved (B) images of nuclei of nontreated (NT) and treated (via NCS and 4-OHT) U2OS and DIvA cells, respectively. Magnified dSTORM images (C) of the selected individual foci. Scale bar: 1 μm (B) and 200 nm (C)

### Spatial distribution of DSB foci within cells

In contrast to traditional optical microscope images, at which the separation of individual foci is a great challenge and their size can only be quantified by their intensity values, cluster-analysis dSTORM images pave the way for quantitative evaluation. Quantitative functions of DSB foci, such as their spatial density variation and their area distribution were evaluated by means of 2D density-based spatial cluster analysis (DBSCAN). This algorithm requires two input parameters: a minimum number of points that forms a cluster (*N*_*core*_) and the maximum distance between two adjacent points (*ε*) ^44^. For the elimination of non-specific labelling and imprecise, out-of-focus localization, *N*_*core*_ and *ε* were set to 8 and 50 nm during the simulations, respectively. Figure 2 A, C and E represent a typical dSTORM super-resolved image of non-treated (NT) and treated (T) (with NCS and 4-OHT) U2OS and DIvA cells. The cluster-analysis images of the selected cells are also shown (Figure 2 B, D and F). In order to efficiently reveal the DSB distribution pattern inside the nucleus, the cluster analysis module was implemented into our rainSTORM localization software (for details see Methods).

**Figure 2.**
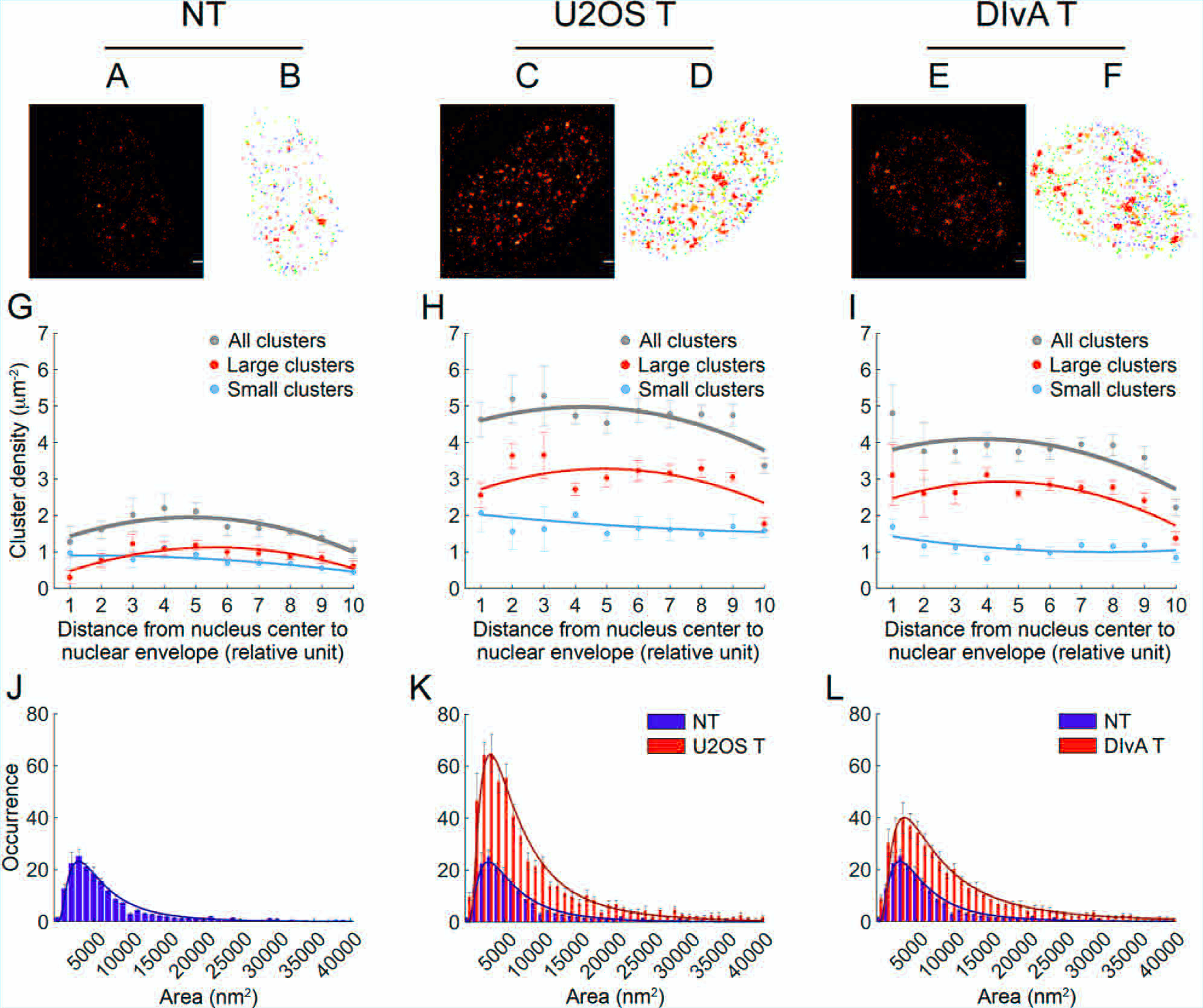
dSTORM (A, C, and E) and cluster-analysis (B, D, and F) images of untreated (A, B) and treated U2OS (C, D) and DIvA (E, F) cells. The average cluster density functions are indicated in grey (G, H and I). The density function calculated for small (<5000 nm2) and large (>5000 nm2) clusters are also shown in blue and red, respectively. Comparative histograms of the area distribution of cluster sizes are presented for untreated (J) NCS treated (K) and DIvA cells (L)

The algorithm isolated and quantified each DSB focus based on their area and spatial distribution inside the nucleus. While Figure 2 A-F show single but typical nuclei, Figure G-L depict the evaluation results of several, 5 untreated, 4 treated U2OS and 6 treated DIvA nuclei, respectively. By using the algorithm, discrete DSB foci were analysed by quantifying the γH2AX tag-pair distances. These defined foci are different from those detected by standard microscopy showing the sub-structures of molecular arrangements (Figure 2 B, D and F).

It was reported that in euchromatic milieu DNA breaks could be repaired more effectively since these breaks do not need to be repositioned outside of the heterochromatic domain for the successful repair^45^. In order to detect the distribution of the γH2AX enriched repair foci within the nucleus we applied cluster recognition, and we compared their position to the centre of the nucleus determined by a computer algorithm. This position was used as a central point to plot individual repair foci by modelling their localization as a circular shell. The spatial density of the clusters shows a non-linear distribution (grey dots and trend lines in Figure 2 G, H and I). Following DSB induction, in comparison to the control nuclei, we could detect almost 3-times more foci formation in the treated U2OS and DIvA cells (grey dots and trend lines in Figure 2 H and I vs G), respectively. The amount of the measured density increases from the periphery towards the centre of the nucleus due to the different chromatin organization. For further quantitative analysis of the clusters, we sorted them into two classes based on their size (indicated in red and blue in Figure 2 G-I). While in the nuclei of untreated cells, both populations have similar and regular density distribution, in the treated cells the number of larger clusters (>5,000 nm^2^) was found to be 2 times more than the number of the small (<5,000 nm^2^) ones (Figure 2 H-I red vs. blue lines). Therefore, the larger clusters could be appeared upon DNA damage induction and it could be differed from the foci induced by endogenous DNA damages.

Finally, for deep evaluation, several individual foci of each treatment category were chosen. For that, the sizes of clusters associated with the DSB foci were categorized by their area, and their distribution was presented in histograms shown on Figure 2 J, K and L. The measured distributions could be fitted with lognormal curves ^37^. In control cells the expected area of the calculated surface was found to be 2,950 nm^2^, and this value was only slightly changed in the treated U2OS and DIvA cells (2,850 nm^2^ and 3,150 nm^2^). However, the mean values of the calculated distributions were increased with 18% (8,750 nm^2^) and 55% (11,550 nm^2^) in the treated cells compared to the untreated ones (7,450 nm^2^), since the normalized occurrence of the large-sized clusters was enriched following DSB induction. The presented data reveal that this algorithm can also be used to separate the γH2AX background, i.e. endogenous versus DSB induced signals (Figure 2 J vs. K and L). These data suggest that DSBs induced by either neocarzinostatin or 4-OHT could result in elevated γH2AX enriched foci both in number and size.

The evaluation of spatial distribution of DSB foci is based on the cluster analysis of the raw dSTORM images, in which the pixel value represents the number of the accepted localizations. However, the number of localizations (*N*_*localizations*_) generated by a single labelled histone strongly depends on the lifetime of the fluorescence ON state (*N*_*lifetime*_), the labelling density (*N*_*labelling*_) and the number of reactivation circles of the applied dye molecules (*N*_*activation*_). Due to multiple localizations, the accepted ones belonging to the same target molecule form a cluster, the size of which depends on the localization precision.

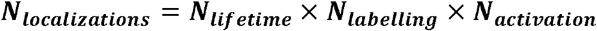

Segmentation and quantitative evaluation of individual blobs are required to determine the response function of dSTORM imaging, in other words the number of localizations belonging to a single labelled histone molecule. Being aware of this response function the size and spatial distribution of the captured foci can be statistically evaluated.

#### Trajectory fitting of individual blinking events (N_lifetime_)

In dSTORM the fluorescence dye molecules are stochastically switched between their OFF (no fluorescence), ON (fluorescence) and bleached states. The occupation of these states can be controlled by a special switching buffer and data acquisition (laser power etc.) parameters ^46^. The lifetime of the ON state strongly depends on the biological sample and the local chemical environment ^47^. Ideally, the lifetime of the ON state is in accordance with the exposure time, and the captured photons emitted by a single dye molecule can be visualized on a single image frame. However, the detector is not triggered, and the lifetime of ON state is not constant. As a result, the same dye molecule can be captured on sequential frames and the trajectory length of a single emitted fluorescence shows an exponential decay (Figure 3A). A trajectory fitting module was built into the rainSTORM localization software that can realign these sections^48^. This resulted in less but more precise detection of localizations (Figure 3 B). Labelling density, buffer condition and image acquisition parameters were set to minimize the possibility of spatial and temporal overlap of individual PSFs (Point Spread Function), hence single Gaussian fitting could be used throughout this work. Figure 3 shows the dSTORM (Figure 3 C and D) and the cluster analysis images (Figure 3 E and F) of a focus before and after trajectory fitting, respectively. The trajectory fitted image reveals more structural details, and consequently provides a more appropriate data source for cluster analysis (Figure 3 D and F).

**Figure 3.**
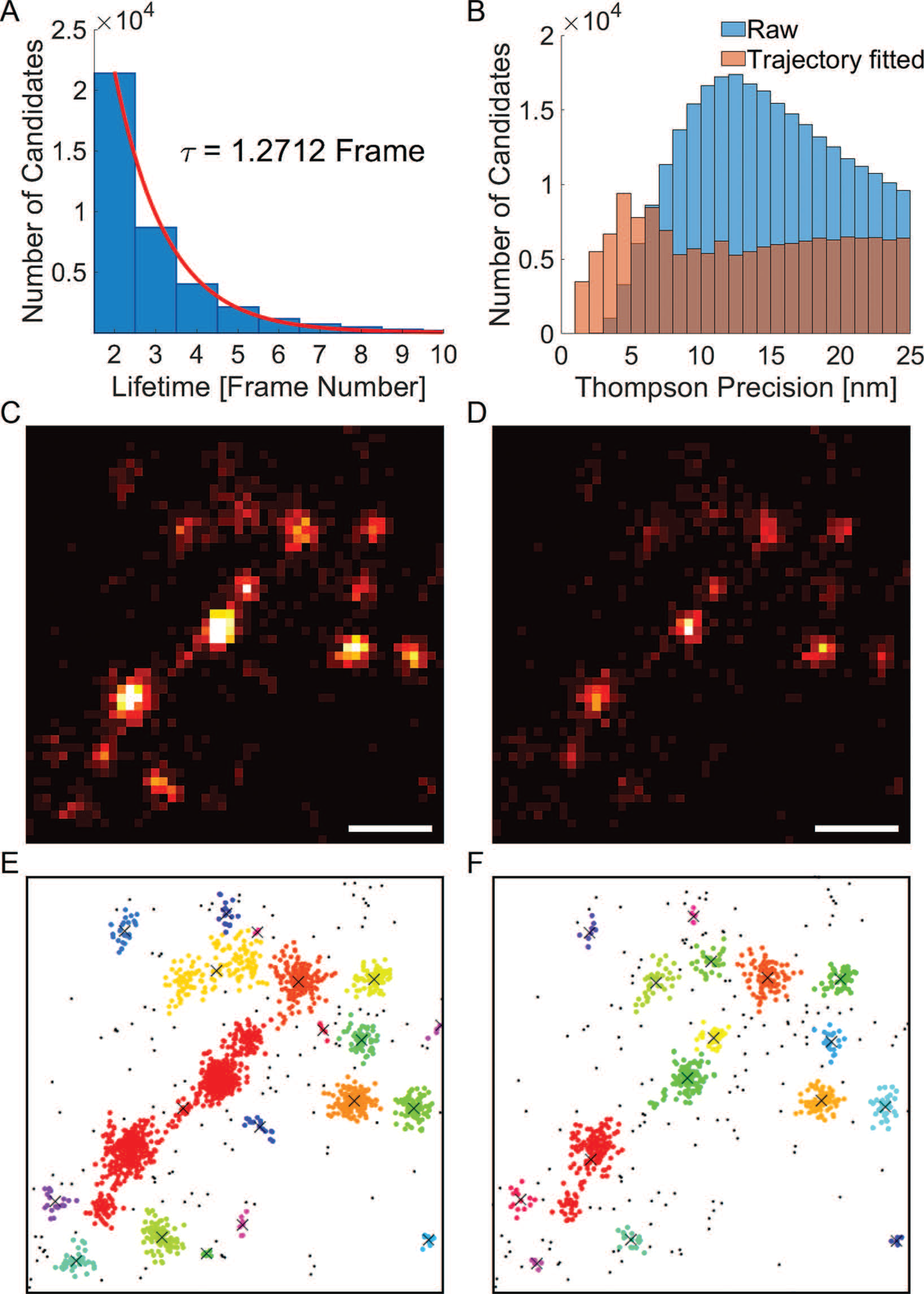
The histogram represents the lifetime of the ON state (A). Trajectory fitting precise localizations (orange in B) and the original distribution (blue in B) are represented in B. 2D original (C) and trajectory fitted dSTORM image (D) of the same focus, respectively. Scale bar is 200 nm. Graphical representation of cluster analysis of a focus before (E) and after (F) trajectory fitting, respectively. The different colours indicate different sub-clusters within a γH2AX cluster.

#### Number of blinking per individual nano-foci (N_labelling_×N_activation_)

Generally, during immunostaining techniques, proteins are recognized by primary and fluorophore-conjugated secondary antibodies (Figure 4 A). However, the number of molecules, taking part in the labelling procedure and then in super-resolution imaging, strongly depends on the actual biological sample (number of epitopes etc.) and the local environment (pH, permeability etc.). During our measurements, due to the sterical hindrance of the nucleosomes, a single primary antibody can bind to the target γH2AX molecule. In addition, our measurements also support already published data that the connection between the primary and secondary antibody is not equal (i.e. IgG), since the 2^nd^ antibody could recognize two epitope surfaces on the first antibody binding ^49^. Consequently, a single γH2AX molecule is labelled by one primary and one or two secondary antibodies. The number of dye molecules per secondary antibody was set to four based on consultations with the manufacturer. In conclusion, we used a model in which a single target histone molecule is labelled either by 4 or 8 fluorescent dye molecules, therefore the number of dye molecules per γH2AX molecule is

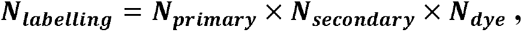

where *N*_*primary*_=1, *N*_*secondary*_=1 or 2 and N_*dye*_=4. The real ratio of γH2AX labelled by 4 or 8 fluorescent dye molecules could be determined by means of cluster analysis (DBSCAN). To eliminate larger clusters belonging to multiple γH2AX foci, *N* and *ε* were set to 5 and 25 nm during the simulations, respectively. After further filtering steps, only the small clusters (area<5000 nm^2^) were evaluated by providing us a high likelihood that all the accepted clusters were associated with the footprint of a single γH2AX nano-focus. These clusters are represented with dark blue colour in Figure 4 D. The histograms of localizations per nano-focus were depicted by using four different image stack sizes and were fitted with a theoretical curve (Figure 4 E). This curve is a linear combination of the two distributions representing the cases of 4 and 8 dyes/γH2AX. Based on the weight of the two components, the ratio of γH2AX molecules labelled with a single or two secondary antibodies can be determined ^49^.

**Figure 4.**
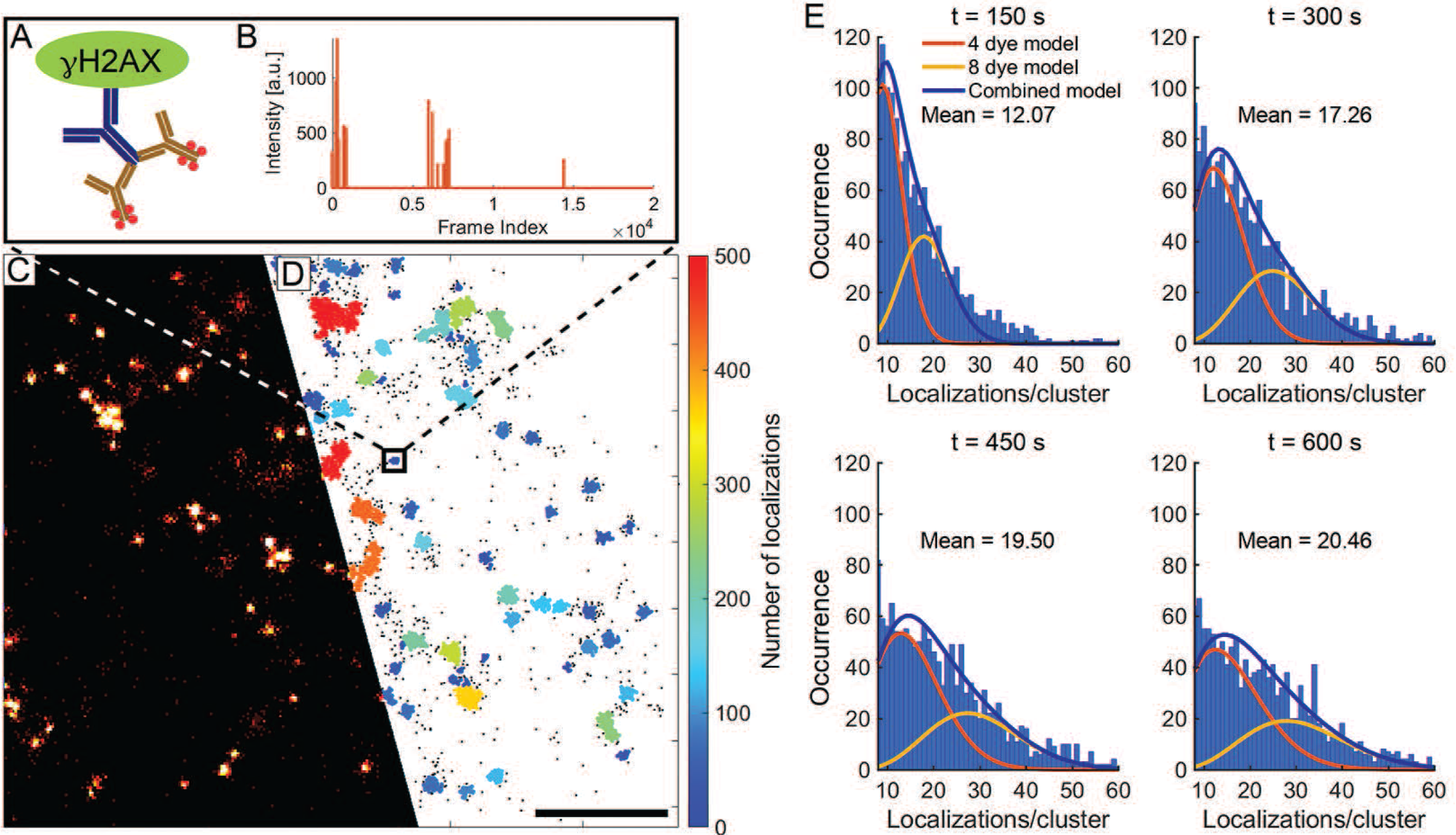
Schematic representation of the binding of first and secondary antibodies to a yH2AX (A) and their frame indexes are shown (B). Super-resolved dSTORM image before (C) and after (D) cluster analysis. Images taken during dSTORM (C) and the clusters containing γH2AX molecules were selected via cluster analysis (D) and their histogram was applied to determine the ratio of labelling via 4 and 8 dye molecules (E) and the response function. Scale bar represents 1 micron.

Additionally, this ratio also depends on the duration of the measurement, since photobleaching plays a central role in the model. In dSTORM technique dye molecules can be switched ON and OFF several times before they are finally bleached. It was shown that in a three-state switching model ^47^ the number of switching circles follows Poisson and geometrical distributions in short (k_bl_t≪1) and long (k_bl_t≫1) data acquisition times, respectively ^47^. However, the typical number of switching circles (*N*_*activation*_) has been already published ^50^, it should be determined more specifically, since it strongly depends on the sample, the buffer conditions and the data acquisition parameters. Based on the evaluation of the fitted curves (Figure 4 E) it can be realized that a measurement time longer than 500 s (>20,000 image frame with 30 ms exposure time) was required for adequate statistical data analysis. This stochiometric evaluation proved that under the measurement conditions detailed above on average 20 localizations belonged to a single γH2AX molecule, i.e. the response function of the system was found to be 20 localizations/target molecule.

### Quantitative analysis of single DBS foci

Based on the statistically given response function we could determine the number of labelled γH2AX in the individual foci both in the untreated and in the treated cells. dSTORM images and their cluster maps of three randomly selected cells with three typical foci are represented in Figure 5 (A-I). In the treated cells, the density of DSB foci was found to be 2.8-and 2.2-times higher compared to the untreated U2OS control cells (Figure 5 D and G vs. A). In those cells, in which DSBs were induced, an increased number of γH2AX localization could be observed within the DSB focus compared to control cells. The histograms obtained from the quantitative measurements are shown on Figure 5 M, N and O. The distribution of cluster sizes based on their γH2AX values (Figure 2 J, K and L) follows the same kinetic. Consequently, the number of γH2AX molecules within a cluster is linearly proportional to its area in untreated (397±7 *γH2AX/nm^2^*), treated U2OS (427±3 *γH2AX/nm*^*2*^), and treated DivA (412±4 *γH2AX/nm*^*2*^) cells shown on Figure 5 J, K and L, respectively.

**Figure 5.**
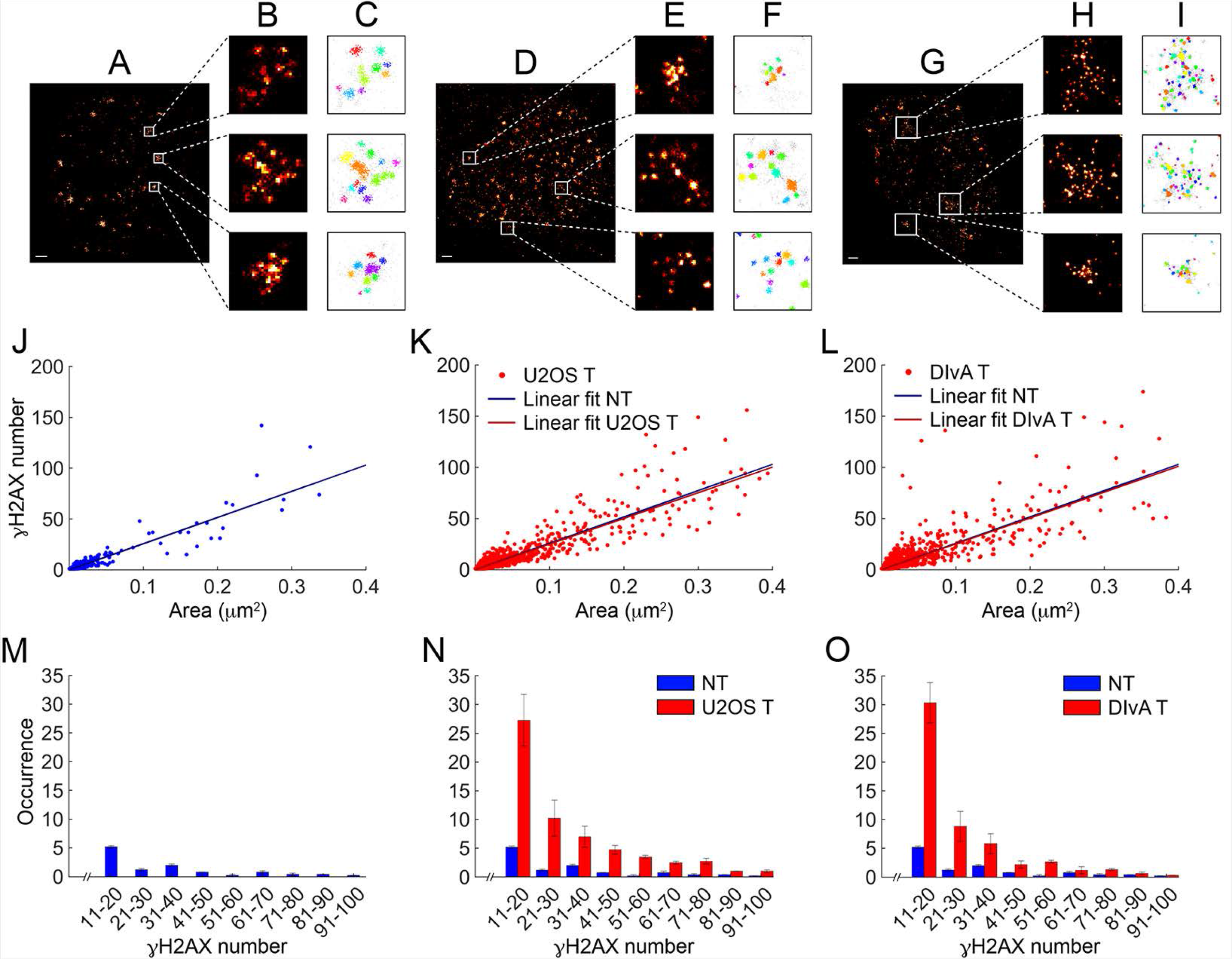
Super-resolved dSTORM images of the entire nuclei of untreated (A) and treated U2OS (D) and DIvA (G) cells. Three typical foci were selected (B, E and H) and cluster analysed (C, F, and I).The number of γH2AX as function of the area of the cluster is depicted and fitted by a linear curve (J, K, and L) based on evaluation of 5 untreated, 4 treated U2OS and 6 treated DIvA cells. Histograms of the γH2AX number/cluster (M, N and O) using the same data show a similar distribution as the area distribution.

The major advantage of our algorithm is that it allows the schematic representation of the individual localization of repair foci and we could apply a topological analysis of the captured images. By using the parameters (localization, primary and secondary antibody number, fluorophores, etc.) determined in our measurements, we could mathematically analyse the topology of DNA repair clusters within a focus. Each blinking event was measured, quantified and following the calculations the number of the independent γH2AX positions were plotted into two-dimensional complexes (Figure 5 A-I). In the representation process each point which localized in proximity (N=8 and *ε*=50 nm) were considered to belong to the same cluster. The described plots of each condition (U2OS control, NCS treated U2OS and 4-OHT treated DIvA cells) are shown in Figure 5 C, 5 F and 5 I. These characterizations allow a compact and illustrative visualization of specific sub-structures. Based on this plot we could tag the point structures with barcodes, which provides novel possibilities to analyse and categorize the number of γH2AX clustered sub-domain structures in cell nuclei (Figures 5 J-L). These representations demonstrate that the endogenous and induced DSBs are covered by approximately 10-50 H2AX S139 phosphorylated histones, which implies an approximately 20-40 kb DNA region (Figures 5 M-O) ^51,52^.

## DISCUSSION

DNA double-strand breaks are one of the most harmful DNA damages since the dsDNA strand loses its integrity and the improper association of these broken DNA strands could lead to chromosomal rearrangements. During DSB repair, the chromatin structure is rearranged and H2AX S139 phosphorylation rapidly appears around the damage sites ^5,6,24^. These steps allow the efficient recruitment of the repair factors to the damaged DNA regions and implicate in the choice between the DNA repair pathways. For examining the DSB-induced chromatin changes, confocal microscopy-based techniques are used in most of the studies, although in the last few years high-throughput chromosome conformation capture technique (4C) and single cell microscopy were utilized to gain detailed insights about the protein interactions and cascades involved in the different repair pathways ^31,53–57^. A more detailed overview has raised more questions, which could be answered only at a single-cell level: how the different DNA repair pathways are chosen and how individual repair proteins are regulated to access the DNA repair site. Answering these questions requires a better resolution, most favourably in a single molecule detection level deeply into the mechanistical organization of the orchestrated repair focus. For a single molecule detection, the 200-300 nm resolution, which is the limitation of the conventional microscopy would not provide sufficiently detailed image resolution. Recently G. Legube’s laboratory has published detailed information about the chromatin organization of DNA repair centres by using 4C^53^. However, the limitation of the technique is that it shows the average of a given focus by combining the data obtained from a large population of cells. Recent applications of electron-microscopy and super-resolution light microscopy ^24,30,31,55–57^ have demonstrated that it is feasible to study single molecular arrangements within a repair focus. By improving the resolution of microscopy and data evaluation of structures in meso-and nano-scale level, the searching for the best-suited analysis parameters and potentially useful classification criteria of repair foci and damaged chromatin sites have become to be important.

In this study we addressed the nano-scale resolution of a single repair focus by quantitative dSTORM technique in order to reveal the structure of γH2AX containing foci within the nuclear environment. For this, we quantified numerous parameters, such as the number of fluorophores, primary and secondary antibodies, which could bind to a single target molecule, etc. and we applied these parameters to evaluate the images by using dSTORM based image processing. By this unconventional procedure we provided 20 nm resolution imaging followed by cluster analysis of various repair loci. However, the quantitative dSTORM technique has been used for studying cellular events, such as cytoskeleton formation in the cytoplasm, in our study it has been utilized for the first time to study cellular events in the nucleus. By the data we obtained from our quantitative measurements it is the first demonstration of the deep structure of a DNA repair focus, at which a single nucleosome resolution has been obtained together with the γH2AX sub-domain cluster organization. Our data suggest a looping mechanism, in which approximately twenty S139 phosphorylated H2AX histones are included within a single chromatin sub-domain, which are localized within an approximately 40-50 kb DNA region^51,52^.

Additionally, another important finding of our study is that a single repair focus contains approximately 10 units of γH2AX enriched sub-cluster. However, we could not determine whether it is a single DSB or several broken DNA regions are associated in one focus. Since γH2AX clusters spatially distribute in the nuclear space according to a pattern that is dependent on the progression of DDR. This pattern recapitulates the previously described repair kinetics, underlying an euchromatin-to-heterochromatin repair trend since it was shown that heterochromatin regions require further structural remodelling before specific DNA repair proteins could assess to those regions. These data highlight another mechanism, in which the complex DNA breaks could be associated in repair centres for efficient DNA repair. This question could be answered in the future by using our quantitative dSTORM method.

In conclusion, we could show that dSTORM is the most adequate tool for deep investigation of DNA double-strand break induced repair focus formation. We believe that nowadays this is the most appropriate procedure for quantitative analyses of the structural changes of a single repair focus in individual cells at nano-scale resolution. The measurements and the procedure we applied in our study allow ultra-resolution insights into structures and architectures, offering new perspectives for further understanding the mechanisms of chromatin function in DNA repair.

## AUTHOR CONTRIBUTIONS

Conceived the project and designed the experiments: T.P., M.E., H.M., D.V., Zs.U., Performed the experiments: H.M., D.V., Analysed the data: H.M., D.V., Zs.U., T.P., M.E. Contributed reagents/ materials/ analysis tools: M.E., T.P. Wrote the paper: T.P., M.E., H.M., D.V., Zs.U.

## MATERIALS AND METHODS

### Trajectory fitting algorithms

The exposure time in localization microscopy is matched to the ON state lifetime of individual molecules. However, due to the stochastic feature of the blinking process, a single fluorescent molecule is typically captured in several sequential frames. Trajectory fitting is an inbuilt algorithm in the rainSTORM localization software that links together photons emitted by the same dye molecule. Localizations on sequential frames which are closer to each other than a preliminary defined *Acceptance Radius* are assumed to belong to the same fluorescence dye molecule. As a result, the code calculates the weighted localization coordinates taking into consideration the captured photon numbers. Therefore, the higher the localization precision, the higher the weight factor ^58^.

### Determination of cluster density

A Matlab code was written to determine the spatial density of clusters inside the nuclei using localization data provided by rainSTORM. First the selected nucleus was segmented with a simple and irregular *N*_*polygon*_-sided (*N*_*polygon*_≈100) polygon using the sum image of the captured frames. The centre of the polygon ^59^ was calculated and connected to all the vertices of the polygon, and all these lines were segmented into ten equal parts (nine division points). In the next step, ten polygons were formed by the n^th^ division points of each line. Clusters inside the i^th^, but outside the (i−1)^th^ polygons were counted and the normalized area cluster density was calculated in each.

### Implementation of 2D/3D DBSCAN into rainSTORM

A DBSCAN based cluster analysis module was implemented into the rainSTORM program. After the reconstruction of the high resolution (SupRes) image the user can select a region using the box tracking tool, and set the two cluster analysis parameters (*N*_*core*_, *ε*). The program plots and saves data for further evaluation and visualization. Larger areas (entire nuclei etc.) can also be selected, but the code automatically segments them into smaller regions to avoid computation fails. After cluster analysis is performed for all sub-regions, the code saves the merged data.

### Experimental determination of bleach rate

The number of cumulative localizations (*N*_*cumulative*_) as a function of time follows an exponential curve the decay of which is proportional to the bleach rate (*k*_*bl*_):

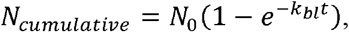

where *N*_*0*_ is the average number of switching cycles of the fluorophore. The two parameters (*k*_*bl*_, *N*_*0*_) were determined by fitting the theoretical curve to the measured data.

### Statistics of *N*_*dye*_ independently switching fluorophores

Fluorescent switching was described by a three-state (ON, OFF and bleached) model. The probability of detecting *n* blinking of *N*_*dye*_ fluorophores is

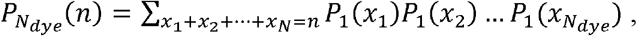

where *x*_*i*_ gives the blinking number of the *i-th* molecule and *P*_*1*_*(m)* is the probability of *m* blinking of a single fluorophore ^47^. Due to the assumption that single γH2AX molecules were labelled by 4 or 8 fluorophores, the overall probability was given as the linear combination of the probabilities

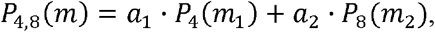

where *m*_*1*_+*m*_*2*_=*m* gives the blinking number, and the ratio of *a*_*1*_ and *a*_*2*_ parameters can be determined by fitting. (see Supplementary Note 1).

### Cell lines, media and culture conditions

DIvA cells were cultured at 37 °C in DMEM (Dulbecco’s Modified Eagle Medium 4.5 g/l glucose, supplemented with L-pyruvate; Lonza,) supplemented with 10 % foetal bovine serum (Lonza), 4 mM L-Glutamine (Sigma-Aldrich), 1 mM puromycin (Gibco) and 1 % antibiotic (Lonza). U2OS osteosarcoma cells were cultured at 37 °C in DMEM (Dulbecco’s Modified Eagle Medium; Lonza) supplemented with 10 % foetal bovine serum (Lonza), 4 mM L-Glutamine (Sigma-Aldrich) and 1 % antibiotic (Sigma-Aldrich).

Both cell lines were grown under standard conditions.

U2OS cell line was purchased from ATCC, DIvA cells were provided by G. Legube. All experimental protocols were approved by the guidelines of the University of Szeged and the Medical Research Council.

### Neocarzinostatin (NCS) treatment

U2OS cells were treated with 5 ng/ml concentration of neocarzinostatin and incubated for 15 minutes. Following the treatment, cells were washed with PBS (phosphate-buffered saline) and incubated in cultured medium for 2 hours.

### 4-hydroxytamoxifen (4-OHT) treatment

DIvA cells were treated with 1 μM concentration of 4-OHT and incubated for 2 hours for the nuclear transport of AsiSI endonuclease. Following the treatment, cells were washed with PBS and then were immunostained.

### Immunocytochemistry

Cells were washed with PBS then incubated with CSK buffer 3 times for 3 minutes and once for half minute [10 mM Hepes pH 7.0 (Sigma-Aldrich), 100 mM sucrose (Sigma-Aldrich), 3 mM MgCl_2_ (Sigma-Aldrich), 0.7 % Triton X-100 (Sigma-Aldrich), 0.3 mg/ml RNase A (Roche)]. Cells were washed twice with PBS, then fixed with 4 % formaldehyde (Sigma-Aldrich) for 10 minutes. Cells were permeabilized with 0.2 % Triton X-100/PBS for 5 minutes. After washing steps, cells were blocked with 5 % BSA (Sigma-Aldrich) in PBST [0.1 % Tween 20 (Sigma-Aldrich) in PBS], supplemented with GAR HRP antibody in 1:200 dilution for 20 minutes. Cells were washed with PBST, then incubated with primary antibodies diluted in 1 % BSA/PBST: anti-γH2AX (Abcam, ab2893) in 1:400 dilution. After washing steps, the following secondary antibody was used: GAR Alexa 647 (Abcam, ab150091) in 1:1500 dilution. After several washing steps with PBST the experiments were conducted after the addition of imaging buffer, which is an aqueous solution diluted in PBS containing an enzymatic oxygen scavenging system GluOx (2,000 U/ml glucose-oxidase (Sigma-Aldrich), 40,000 U/ml catalase (Sigma-Aldrich), 25 mM potassium chloride (Sigma-Aldrich), 22 mM tris(hydroxymethyl)aminomethane (Sigma-Aldrich), 4 mM tris(2-carboxyethyl)phosphine (TCEP) (Sigma-Aldrich)) with 4 % (w/v) glucose (Sigma-Aldrich) and 100 mM β-mercaptoethylamine (MEA) (Sigma-Aldrich). The final pH was set to 6.0-8.5.^60–62^

### dSTORM microscopy

We used a Nikon Eclipse Ti-E frame with a Nikon CFI Apochromat TIRF objective (NA 1.49, 100× magnification, oil immersion) for imaging. EPI-fluorescent illumination was applied at excitation wavelength of 647 nm (2RU-VFL-P-300-647-B1, 300 mW, MPB Communications Ltd.). A filter set from Semrock was used in the microscope (Di03-R405/488/561/635-t1-25×36BrightLine® quad-edge quad-edge super-resolution / TIRF dichroic beamsplitter and FF01-446/523/600/677-25BrightLine® quad-band bandpass filter). An Andor iXon3 DU897 EMCCD camera was used for image acquisition (pixel size: 16 μm) with the following acquisition parameters: 30 ms exposure time, EM gain of 100, temperature of −75 °C

## SUPPLEMENTARY INFORMATIONS

### Supplementary Note 1: Switching statistics of multiple labelling

In a three-state switching model, the time-dependent probability of *m* switching circles of a single fluorescent dye molecule is [2016_Nieuwenhuizen]:

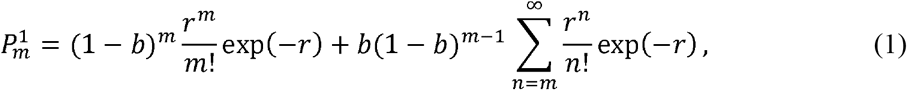

where parameters *r* and *b* depend on the *k*_*sw*_ switching and *k*_*bl*_ effective bleaching rates as *r*=*k*_*sw*_*t* and *b*=*k*_*bl*_/*k*_*sw*_.

In practice, using immunohistochemical procedures, several fluorescence dye molecules label the target molecule and their common switching pattern provides the detected signal. The number of fluorescence dye molecules depends on the stoichiometry of the labelling. Therefore, it is essential to determine the overall probability of *m* switching circles of *N* independent dye molecules (*P*_*m*_^*N*^). It can be given as the sum of probabilities of all the possible cases when *N* molecules generate *m* switching circles:

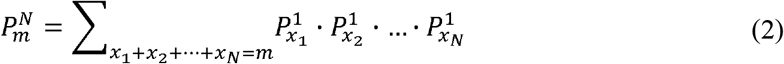

where *x*_*1*_, *x*_*2*_, and *x*_*N*_ mark the number of switching circles can be associated to the 1^st^, 2^nd^ and n^th^ dye molecules, respectively.

As an example, let us assume that only 2 independent dye molecules label the target molecule and provide *m* switching circles. In other words, the total number of switching circles is *m* but we do not know how many switching circles belong to each dye molecules. If the first one was detected *x*_*1*_ time, the second one must be detected x_2_=*m−x*_*1*_ times and the overall probability can be given as the sum of all the possible cases 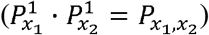:

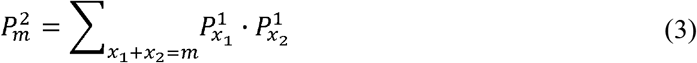

In general, all the possible cases can be calculated and can be arranged in a matrix form. In this representation the sum of elements of the m^th^ minor diagonal gives the overall probability of *m* switching circles generated by two dye molecules. It can be shown that after a critical cluster size the larger that matrix (the larger the possible number of switching circles), the smaller the sum of the minor diagonal elements (smaller the probability of the effective switching circles). The sums of the minor diagonal elements form a vector and give the probability distribution of the switching circles.

**Table S1:**
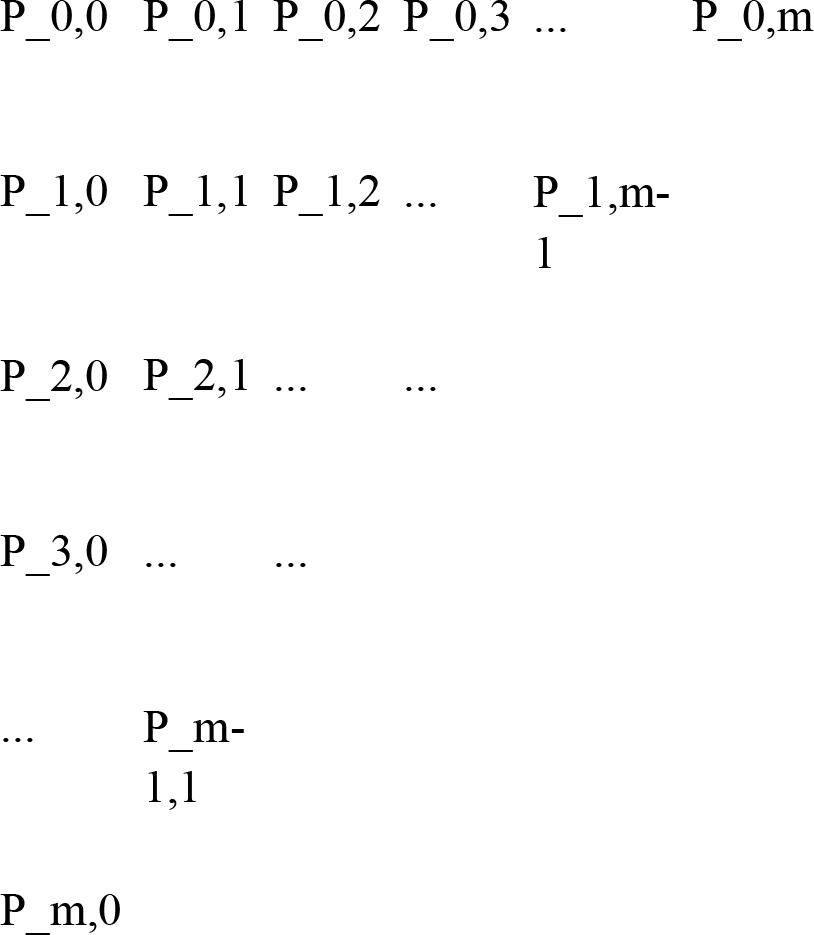
Probabilities of all the possible switching cases are arranged in a matrix form.

The method can be generalized further and the probability of *m* switching circles generated by *N* molecules can be calculated.

It is worth to note that the calculation can be simplified by dividing the *N* number of dye molecules into two independent but known populations (e.g. *K* and *N−K*) with number of switching circles of *i* and *m−i*, respectively.

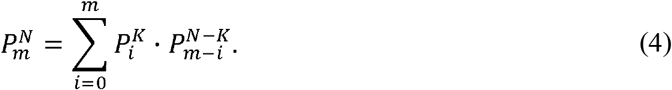

### Supplementary Note 2: Validation of 2D analysis

In this paper all the results and conclusions are based on the evaluation of 2D dSTORM measurements. However, the DSB foci inside the nucleus have a 3D spatial distribution. Therefore, the applicability of the used 2D data analysis requires a validation process. A test sample simulator software (TestSTORM ^63,64^) was used to generate the ground truth model and comparative 2D and 3D evaluations were performed on the reconstructed super-resolved images. Images were evaluated from the same aspect (number of clusters, mean number of cluster elements etc.) as they were studied in the main text.

**Figure S1.**
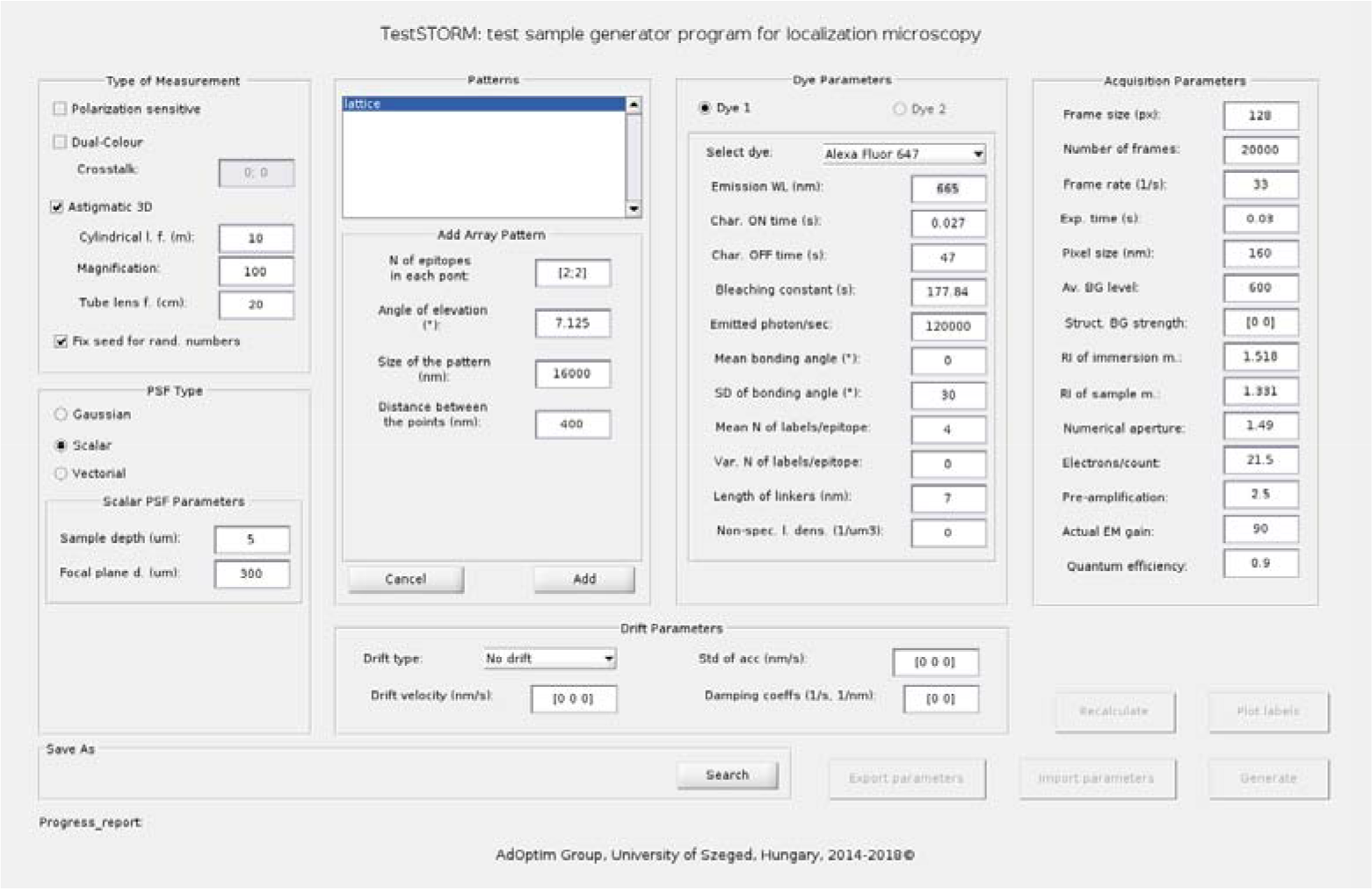
Working GUI window of testSTORM with the used parameters

**Simulation parameters** were matched to the experimental parameters. The most important parameters are depicted in Figure S1, which shows the working GUI window of the TestSTORM code.

A tilted array pattern (lattice) was defined with the following parameters:

Depth inside the sample: Depth_sample_ =5 μm
Refractive index of the sample: n_sample_ = 1.331
Axial range of the sample: Z_range_: (−1 μm,+1 μm)
Axial steps between the adjacent rows: Z_step_ = 50 nm
Distance between the elements (cluster) of the lattice: d = 400 nm
Number of elements in a single column and row: N_cluster_/Z_plane_ = 41
Number of dye molecules per cluster: 8 dye molecules/cluster
Length of the linker: 7 nm

**Figure S2.**
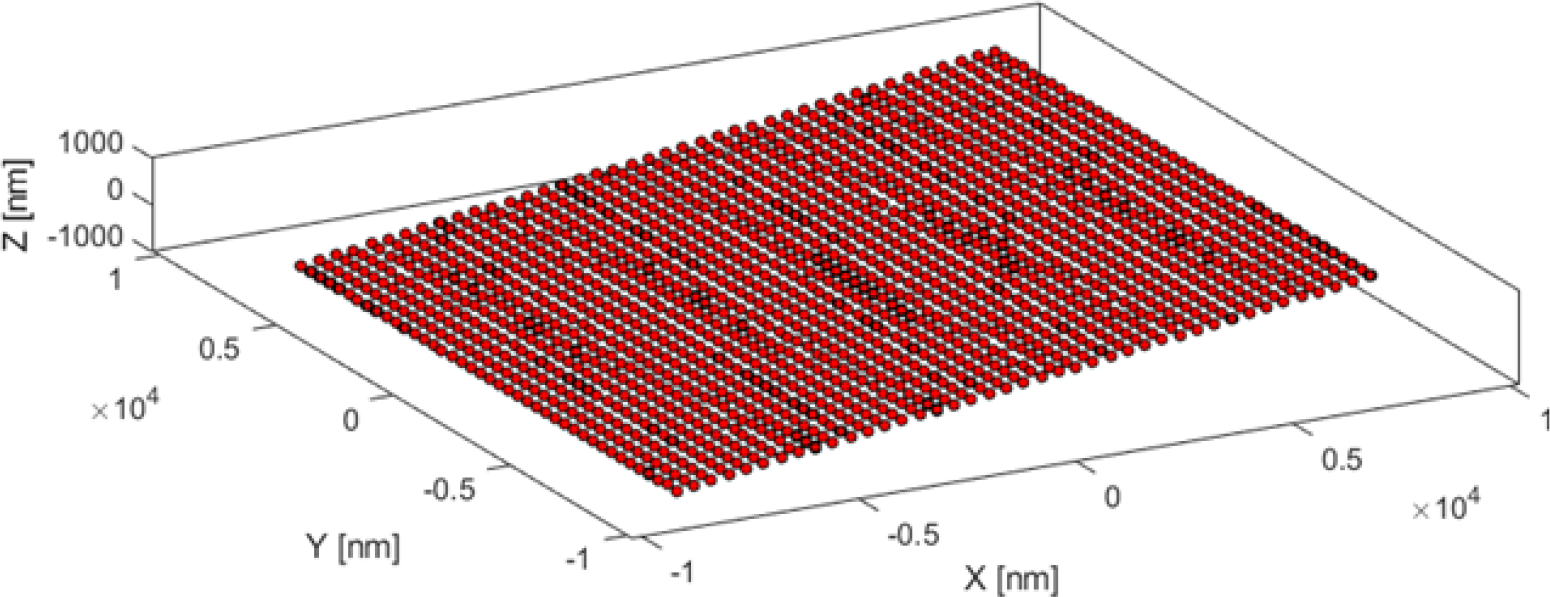
Tilted 3D array pattern consists of 41×41 clusters

A scalar model based on the Pankajakshan-Gibson-Lanni model [2009_Pankajakshan] was applied to calculate the PSF. During the 3D simulations, an additional cylindrical lens with a focal length of 10 m was added to the optical system.

**Figure S3.**
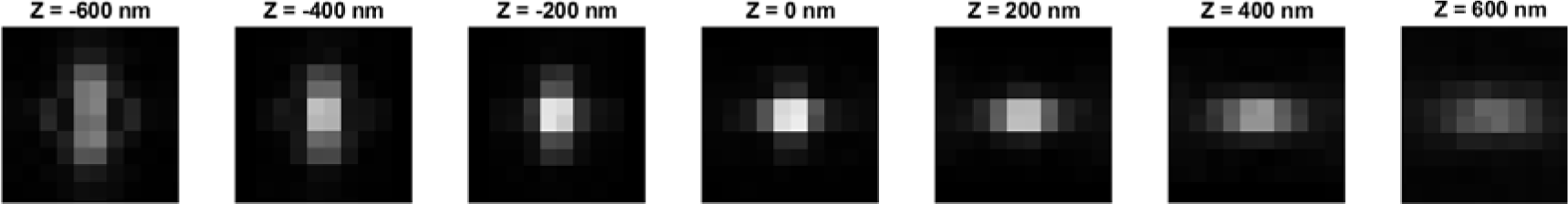
Defocused astigmatic PSF introduced by a cylindrical lens with a focal length of 10 m

High resolution localization images were reconstructed and analysed via the rainSTORM code ^65^ with the following key parameters:

Thompson precision limit: 25 nm
Applied acceptance radius during the trajectory fitting: r_acceptance_ = 50 nm
Residue threshold: 0.06
Lateral cluster analysis distance parameter: ε_xy_ = 50 nm
Axial cluster analysis distance parameter: ε_z_ = 100 nm
Minimum number of points in a single cluster: N_core_=8

**Our simulation results** prove that 2D and 3D imaging provides identical DOF ranges, i.e. dye molecules in the same axial range (~1 μm) can be associated with the accepted localizations (see Figure S4-a). The slightly reduced number of identified clusters in the 3D case (~7%) is caused by the asymmetry of the PSF. This difference does not affect the trend of the evaluation but shows that 3D analysis requires a different calibration process. The mean number of cluster elements (see Figure S4-b) shows an approx. 20% reduction in the 3D case in contrast to the 2D one, and the simulations reveal a slight axial dependence in both cases. During the evaluation this axial dependence was neglected, and an average value was applied. Based on these simulation results one can state that 2D measurements (presented in the main text of the paper) provide reliable data and results for the quantitative evaluations. However, determination of 3D specific merit functions (volume of foci etc.) and features (structure of foci etc.) requires 3D STORM imaging.

**Figure S4.**
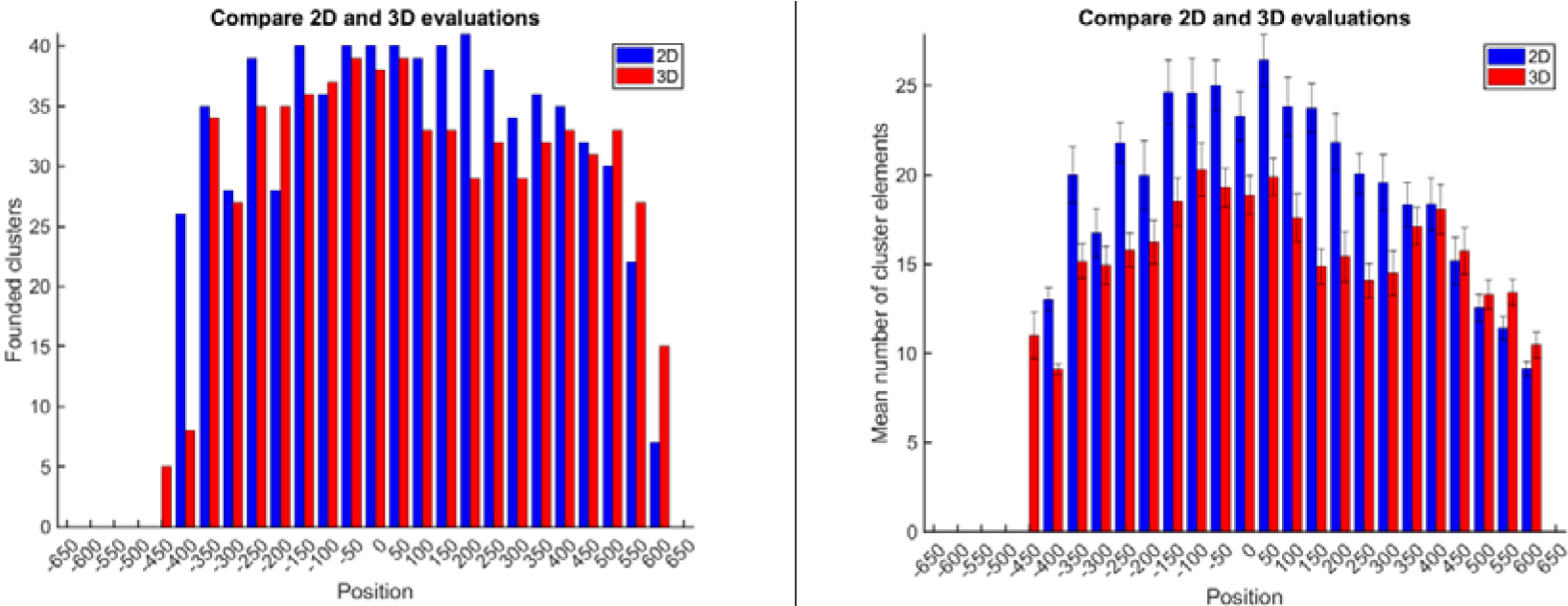
Number of isolated clusters (a) and the mean number of cluster elements as a function of the axial position

